# The feedstock microbiome selectively steers process stability during the anaerobic digestion of waste activated sludge

**DOI:** 10.1101/757922

**Authors:** Cindy Ka Y Law, Rens De Henau, Jo De Vrieze

## Abstract

Strategies to enhance process performance of anaerobic digestion remain of key importance to achieve further spreading of this technology for integrated resource recovery from organic waste streams. Continuous inoculation of the microbial community in the digester *via* the feedstock could be such a cost-effective strategy. Here, anaerobic digestion of fresh waste activated sludge (WAS) was compared with sterilized WAS in response to two common process disturbances, *i.e.*, organic overloading and increasing levels of salts, to determine the importance of feedstock inoculation. A pulse in the organic loading rate severely impacted process performance of the digesters fed sterile WAS, with a 92 ± 45 % decrease in methane production, compared to a 42 ± 31 % increase in the digesters fed fresh WAS, relative to methane production before the pulse. Increasing salt pulses did not show a clear difference in process performance between the digesters fed fresh and sterile WAS, and process recovery was obtained even at the highest salt pulse of 25 g Na^+^ L^−1^. Feedstock sterilisation strongly impacted the microbial community in the digesters. In conclusion, feedstock inoculation can be considered a cheap, yet, disturbance-specific strategy to enhance process stability in full-scale anaerobic digestion processes.

## 1. Introduction

The increasing environmental pollution and energy insecurity are pressing issues in our present society, which makes it important to look for integrated strategies that provide a solution to both issues. The fossil resources around the world are being depleted at a tremendous velocity, *i.e.*, reaching a global total energy use of 1.64 × 10^5^ TWh in 2017 to which renewable sources only contributed 25% (Enerdate 2018). This makes a transition towards sustainable resources for materials and energy of key importance to halt the increase in CO_2_ equivalents emissions and related climate change (De Meester et al. 2012; Hagos et al. 2017). Anaerobic digestion (AD) is a microbial process that can be one of the possible solutions for this problem. The number of full-scale AD plants is still increasing nowadays, even though it already exists for decades (Charnier et al. 2016). The success of AD lies in the fact that it does not only allow the stabilisation of organic waste streams, but it is also a key technology for the recovery of energy (Demitry 2016). The methanogenic archaea are the most critical microorganisms in the AD process, because they are responsible for the production of the energy-dense CH_4_, which can be used for electricity and heat production via a combined heat and power unit (Holm-Nielsen et al. 2009). However, these methanogens are most sensitive to suboptimal process conditions, *e.g.*, the presence of potential toxic compounds and organic overloading, and, hence, are susceptible to process failure (Appels et al. 2008). This sensitivity can in some cases prevent implantation of this technology (Demitry 2016).

One of the most common solution for preventing process failure is anaerobic co-digestion. Co-digestion can improve process stability, by (1) diluting potential inhibitory substances, such as ammonia toxicity, and (2) optimising the nutrient balance (Mata-Alvarez et al. 2011; Mata-Alvarez et al. 2000). Nevertheless, process stabilization and optimization remains difficult (Hubenov et al. 2015; Kacprzak et al. 2010), and the unbalanced availability of suitable feedstocks is another problem that hampers an efficient co-digestion process (Hagos et al. 2017).

Another strategy to improve process stability is bioaugmentation, whereby a specific consortium, either enriched or isolated from similar systems or obtained from other ecosystems, is added to enhance the desired activity (De Vrieze and Verstraete 2016; Schauer-Gimenez et al. 2010). These microorganisms can improve the start-up of new digesters (Saravanane et al. 2001a; Saravanane et al. 2001b), reduce odour emissions (Duran et al. 2006; Tepe et al. 2008), and/or facilitate recovery of the reactors after an organic overload (Lynch et al. 1987). The disadvantage of this method is that it requires a certain volume of the biomass itself to be replaced, *i.e.*, 10 % or more, whereby it is often not cost-efficient (De Vrieze and Verstraete 2016; Fotidis et al. 2014). Usually, only a temporal increase in CH_4_ production can be observed, due to wash-out of the bioaugmented microorganism and/or possible competition with the indigenous microorganisms (Schauer-Gimenez et al. 2010).

The addition of cations is also an alternative to solve the problem of process stability. The influent often contains a suboptimal ratio of the most common cations, *i.e.*, Ca^2+^, Na^+^, K^+^ and Mg^2+^ (Kugelman and McCarty 1965). An imbalance of this ratio can cause inhibition of the methanogens, which leads to process failure. To prevent this occurrence, the addition of other cations can restore the optimal balance in the feedstock, and this should result in optimal conditions for the microorganisms (Appels et al. 2008; Kugelman and McCarty 1965). The downside of this approach is that these cations are costly, *i.e.,* the bulk market price of CaCl2 is around € 150-250 ton^−1^, while the bulk market price of MgCl_2_ ranges between € 250-300 ton^−1^ (www.icis.com, consulted June 2019), and their addition to the feedstock will result in an increase in the conductivity, which can have an overall negative impact on the microbial community. Hence, preventing AD process failure *via* an economically feasible approach is a key aspect that requires further investigation before such a prevention strategy can be applied at the full scale. Such a strategy will further solidify the use of AD, which will result in a higher sustainability in the use and recovery of the energy, combined with an environmental friendly way to treat organic waste streams (Chen et al. 2008).

The aim of this research was to tackle process failure in a cost-efficient manner by considering the importance of continuous inoculation or bio-augmentation through microorganisms in the feedstock. Two different process disturbances, *i.e.*, organic overloading and increasing levels of salts were considered, given the potential different impact of feedstock inoculation. We hypothesized that the microorganisms present in the feedstock can (1) support processes resistance against disturbances, and (2) enhance recovery following process inhibition.

## 2. Experimental procedures

### 2.1. Inoculum and feedstock

The inoculum for the operation of the lab-scale anaerobic digesters was obtained from the full-scale sludge digester of the wastewater treatment plant the Ossemeersen (Ghent, Belgium) (Table 1). The waste activated sludge (WAS) that was used as feedstock was also obtained from the Ossemeersen in two separate batches that were used in the first and second stage of the experiment, respectively (Table 2). The WAS was stored at 4°C until use.

**Table 1.** Characteristics of the inoculum sludge (n=3). TSS = total suspended solids, VSS = volatile suspended solids, COD = chemical oxygen demand, TAN = total ammonia nitrogen, VFA = volatile fatty acids, FA = free ammonia nitrogen.

| Parameter | Unit | Inoculum |
| --- | --- | --- |
| pH | - | 7.57 ± 0.03 |
| TSS | g TSS L<sup>-1</sup> | 50.1 ± 0.1 |
| VSS | g VSS L<sup>-1</sup> | 26.6 ± 0.2 |
| Conductivity | mS cm<sup>-1</sup> | 8.13 ± 0.11 |
| Total VFA | mg COD L<sup>-1</sup> | 0 ± 0 |
| TAN | mg N L<sup>-1</sup> | 1147 ± 23 |
| Na<sup>+</sup> | mg L<sup>-1</sup> | 156 ± 22 |
| K<sup>+</sup> | mg L<sup>-1</sup> | 219 ± 20 |
| FA<sup>1</sup> | mg N L<sup>-1</sup> | 43 ± 1 |

<sup>1</sup>The free ammonia (FA) content was calculated based on the TAN concentration, pH and temperature in the full-scale installation (Anthonisen et al. 1976).

**Table 2.** Characteristics of the two batches of waste activated sludge (n=3). TS = total solids, VS = volatile solids, COD = chemical oxygen demand, VFA = volatile fatty acids, TAN = total ammonia nitrogen, TKN = Kjeldahl nitrogen, FW = fresh weight.

| Parameter | Unit | WAS1 | WAS2 |
| --- | --- | --- | --- |
| pH | - | 7.04 ± 0.01 | 6.37 ± 0.02 |
| TS | g TS kg<sup>-1</sup> FW | 76.2 ± 0.9 | 54.3 ± 0.1 |
| VS | g VS kg<sup>-1</sup> FW | 44.2 ± 0.7 | 33.6 ± 0.1 |
| Total COD | g COD kg<sup>-1</sup> FW | 74.4 ± 9.3 | 44.7 ± 1.1 |
| Conductivity | mS cm<sup>-1</sup> | 4.69 ± 0.01 | 4.30 ± 0.01 |
| Total VFA | g COD kg<sup>-1</sup> FW | 0 ± 0 | 0 ± 0 |
| TAN | mg N kg<sup>-1</sup> FW | 345 ± 13 | 571 ± 7 |
| Na<sup>+</sup> | mg Na<sup>+</sup> kg<sup>-1</sup> | 173 ± 44 | 67 ± 17 |
| K<sup>+</sup> | mg K<sup>+</sup> kg<sup>-1</sup> | 164 ± 1 | 183 ± 3 |
| TKN | mg N kg<sup>-1</sup> FW | 4121 ± 630 | 3321 ± 247 |
| COD:N ratio | - | 18.1 ± 3.6 | 13.5 ± 1.1 |
| TS:VS ratio | - | 1.72 ± 0.03 | 1.62 ± 0.01 |
| COD:VS ratio | - | 1.68 ± 0.21 | 1.33 ± 0.03 |

### 2.2. Experimental design and operation

Six glass Schott bottles with a total volume of 1 L and a working volume of 800 mL were operated as lab-scale anaerobic digesters. The digesters were sealed with air-tight rubber stoppers, and connected to a water displacement system *via* gas-tight PVC tubing to monitor biogas production. The liquid in the water displacement system had a pH < 4.3 to avoid the CO2 in the biogas from dissolving. Gas samples were collected *via* a Laboport® vacuum pump (KNF Group International, Aartselaar, Belgium) and glass sampling tube of 250 mL (Glasgerätebau Ochs, Lenglern, Germany) for further analysis.

The digesters were operated in a semi-continuous stirred tank reactor mode, at mesophilic conditions in a temperature-controlled room at 34 ± 1°C. The reactors were initially filled to a total volume of 800 mL with inoculum, which was diluted with tap water to obtain an initial VSS (volatile suspended solids) concentration of 10 g L^−1^. The digesters were fed manually by briefly opening the digesters, three times a week with WAS. Three digesters (biological replicates) were fed fresh WAS, while the three other digesters (also biological replicates) were fed WAS that was autoclaved (30 min at 121 °C) twice (Table S1). This double autoclaving step, with an incubation period of 6-12 hours at room temperature between the two autoclavation steps, was included to ensure complete sterilisation of the sludge, including spore-forming bacteria.

During the start-up phase (day 0-26), the organic loading rate (OLR) was slowly increased from 0.94 to 3.75 g COD L^−1^ d^−1^ (chemical oxygen demand), and the hydraulic retention time was decreased from 40 to 10 days (Table 3). From day 27-126 on, an OLR of 3.75 g COD L^−1^ d^−1^ was maintained (WAS1), and between day 127-207, an OLR of 4.47 g COD L^−1^ d^−1^ was used (WAS2). The WAS1 was diluted with tap water in a 1:1 ratio to avoid overloading, while the WAS2 was used as such, because of the lower VS content. Glycerol was added on day 48 as a single additional pulse of 1 g COD L^−^ d^−1^ to provoke organic overloading. Different pulses of NaCl were added on day 118 (6.25 g Na^+^ L^−1^), day 160 (12.5 g Na^+^ L^−1^) and day 188 (25 g Na^+^ L^−1^).

**Table 3.** Overview of different phases of the experiment. OLR = Organic loading rate. HRT = Hydraulic retention time. During Phase 1 the first batch of waste activated sludge (WAS1) was used, while during Phase 2 the second batch of waste active sludge (WAS2) was used, hence, the difference in OLR.

| Phase | Period | OLR (g COD L<sup>-1</sup> d<sup>-1</sup>) | HRT (days) |
| --- | --- | --- | --- |
| Start-up | Day 0-12 | 0.94 | 40 |
|  | Day 13-26 | 1.88 | 20 |
| Phase 1 | Day 27-126 | 3.75 | 10 |
| Phase 2 | Day 126-207 | 4.47 | 10 |

The biogas production and composition were monitored three times a week, together with the pH and conductivity of each digester. Biogas production values were reported at standard temperature (273 K) and pressure (101325 Pa) conditions (STP). The sulphate, phosphate, sodium, total ammonium and volatile fatty acids (VFA) concentrations were measured on weekly basis. Samples for microbial community analysis were taken on day 0 (inoculum and WAS), 48, 118, 160, 188 and 207 from each digester, and stored at −20°C until DNA extraction was performed.

### 2.3. Microbial community analysis

The DNA extraction was carried out with the ZymoBIOMICS™ DNA Miniprep Kit (Zymo Research, Irvine, CA, USA), using a PowerLyzer® 24 Bench Top Bead-Based Homogenizer (MO BIO Laboratories, Inc, Carlsbad, CA, USA), in accordance with the instructions of the manufacturer. Agarose gel electrophoresis and PCR analysis were used to determine the quality of the DNA extracts. The PCR was carried out using the bacterial primers 341F (5’-CCTACGGGNGGCWGCAG) and 785Rmod (5’- GACTACHVGGGTATCTAAKCC), targeting the V3-V4 region of the 16S rRNA gene, following an in-house PCR protocol (Boon et al. 2002; Klindworth et al. 2013). After quality validation, the DNA extracts were sent to BaseClear B.V., Leiden, The Netherlands, for Illumina amplicon sequencing of the bacterial community, using the abovementioned primers, on the MiSeq platform with V3 chemistry. Amplicon sequencing and data processing are described in detail in SI (S2). Real-time PCR analysis was carried out to quantify total bacteria, the methanogenic orders *Methanobacteriales* and *Methanomicrobiales*, and the methanogenic families *Methanosaetaceae* and *Methanosarcinaceae*, as described in SI (S3).

### 2.4. Statistical analyses

Following the data processing of the amplicon sequencing data, a table was generated with the relative abundances of the different OTUs (operational taxonomic units) and their taxonomic assignment (Supplementary file 2) of each sample. Statistical analyses were carried out in R version 3.3.1 (http://www.r-project.org) (R Development Core Team 2013). The similarity of the bacterial community in biological replicates was statistically validated (*P* < 0.05) using a repeated measures analysis of variance (ANOVA, *aov* function) (Connelly et al. 2017). Rescaling of the samples was carried out *via* the “common-scale” approach, by taking the proportions of all OTUs, multiplying them with the minimum sample size, and rounding them to the nearest integer (McMurdie and Holmes 2014). Rarefaction curves (Figure S1) were generated to estimate the degree of “coverage” of the bacterial community (Sanders 1968). The R packages vegan (Oksanen et al. 2016) and phyloseq (McMurdie and Holmes 2013) were used for bacterial community analysis.

Heatmaps were created at the Phylum, Order, Class, Family, and OTU level using the *pheatmap* function (pheatmap package) for which the biological replicates were collated as described earlier (Connelly et al. 2017). Significant differences in microbial community composition between the digesters fed fresh and sterile WAS were identified by means of pair-wise Permutational ANOVA (PERMANOVA) with Bonferroni correction, using the *adonis* function (vegan). The order based Hill’s numbers were used to evaluate the α-diversity in the different digesters. These Hill’s numbers represent richness (number of OTUs, H_0_), the exponential of the Shannon diversity index (H_1_) and the Inverse Simpson index (H_2_). Significant differences in α-diversity between the digesters fed fresh and sterile WAS were determined with the Kruskal-Wallis rank sum test (*kruskal.test* function) (Hill 1973). Correlations between the sodium concentration and Hill’s numbers were determined using the Kendall’s tau correlation (*cor.test* function). The OTUs with a significant difference (*P* < 0.05) in relative abundance between the digesters with fresh or sterilized WAS as feedstock were determined with the *DESeqDataSetFromMatrix* function from the DESeq2 package (Love et al. 2014).

### 2.5. Analytical techniques

Total solids (TS), total suspended solids (TSS), volatile suspended solids (VSS), volatile solids (VS), Kjeldahl nitrogen (TKN) and total COD were measured according to Standard Methods (Greenberg et al. 1992). Soluble COD was measured using Nanocolor COD1500 or 15000 test kits (Machery-Nagel, Düren, Germany), according to the manufacturer’s instructions. The concentrations of NH_4_+, Na^+^ and K^+^ were measured on a 761 Compact Ion Chromatograph (Metrohm, Herisau, Switzerland), which was equipped with a Metrosep C6-250/4.0 main column, a Metrosep C4 Guard/4.0 guard column and a conductivity detector. The eluent contained 1.7 mM HNO3 and 1.7 mM dipicolinic acid. Samples were centrifuged at 3000*g* for 3 min with a Labofuge 400 Heraeus centrifuge (Thermo Fisher Scientific Inc, Merelbeke, Belgium), filtered over a 0.22 µm filter (type PA-45/25, Macherey-Nagel, Germany) and diluted with Milli-Q water to reach the desired concentration range for quantification between 1 and 100 mg L^−1^. The pH was measured with a C532 pH meter, and conductivity was determined with a C833 conductivity meter (Consort, Turnhout, Belgium). The biogas composition was measured using a Compact Gas Chromatograph (Global Analyser Solutions, Breda, The Netherlands) (S5). The different VFA (C2-C8) were measured with a GC-2014 Gas Chromatograph (Shimadzu®, The Netherlands) (S6).

### 2.6. Data submission

The raw fastq files that served as a basis for the bacterial community analysis were deposited in the National Center for Biotechnology Information (NCBI) database (Accession number PRJNA540741).

## 3. Results

### 3.1. Digester performance

#### 3.1.1. The impact of an organic shock load

The initial start-up period during which the OLR was steadily increased and hydraulic retention time decreased (Table 3) over the first 27 days showed a steady increase in performance with increasing methane production rates (Figure 1). For the next 21 days, steady-state conditions were obtained in both digesters from day 27-47 (Figure 1), reflected in a stable methane production of 157 ± 33 and 316 ± 30 mL CH_4_ L^−1^ d^−1^ in the digesters fed fresh and sterile WAS, respectively. The addition of glycerol caused a differential effect in the digesters fed fresh and sterile WAS. A methane production rate of 224 ± 10 mL CH_4_ L^−1^ d^−1^ was observed in the digesters fed fresh WAS on day 53, which is a 42 ± 31 % increase compared to the previous 21 days, because of the additional carbon source. In contrast, methane production decreased to only 25 ± 12 mL CH_4_ L^−1^ d^−1^ in the digesters fed sterile WAS, reflecting a 92 ± 45 % decrease compared to the previous 21 days. This was also reflected in a pH value of 7.16 ± 0.03 for the digesters fed fresh WAS, while the pH in the digesters fed sterile WAS decreased to 6.68 ± 0.00. The recovery period was much longer for the digesters fed fresh WAS in comparison with the digesters fed sterile WAS. Methane production and pH in the digesters fed sterile WAS reached the same values as prior to the glycerol shock only on day 71, while for the digesters fed fresh WAS, complete recovery was already the case on day 53 (Figure 1 & 2). The residual VFA concentration also showed an increase following the glycerol pulse (Figure 2b). The increase in VFA production was a factor two higher for the digesters fed sterile WAS than for the digesters fed fresh WAS, *i.e.*, maximum concentrations of 4.15 ± 0.50 g COD L^−1^ (day 57) and 1.96 ± 0.13 g COD L^−1^ (day 50), respectively, were obtained (Figure 2b). Overall, more time was needed for the recovery of the digesters fed sterile WAS than for the digesters fed fresh WAS.

**Figure 1.**
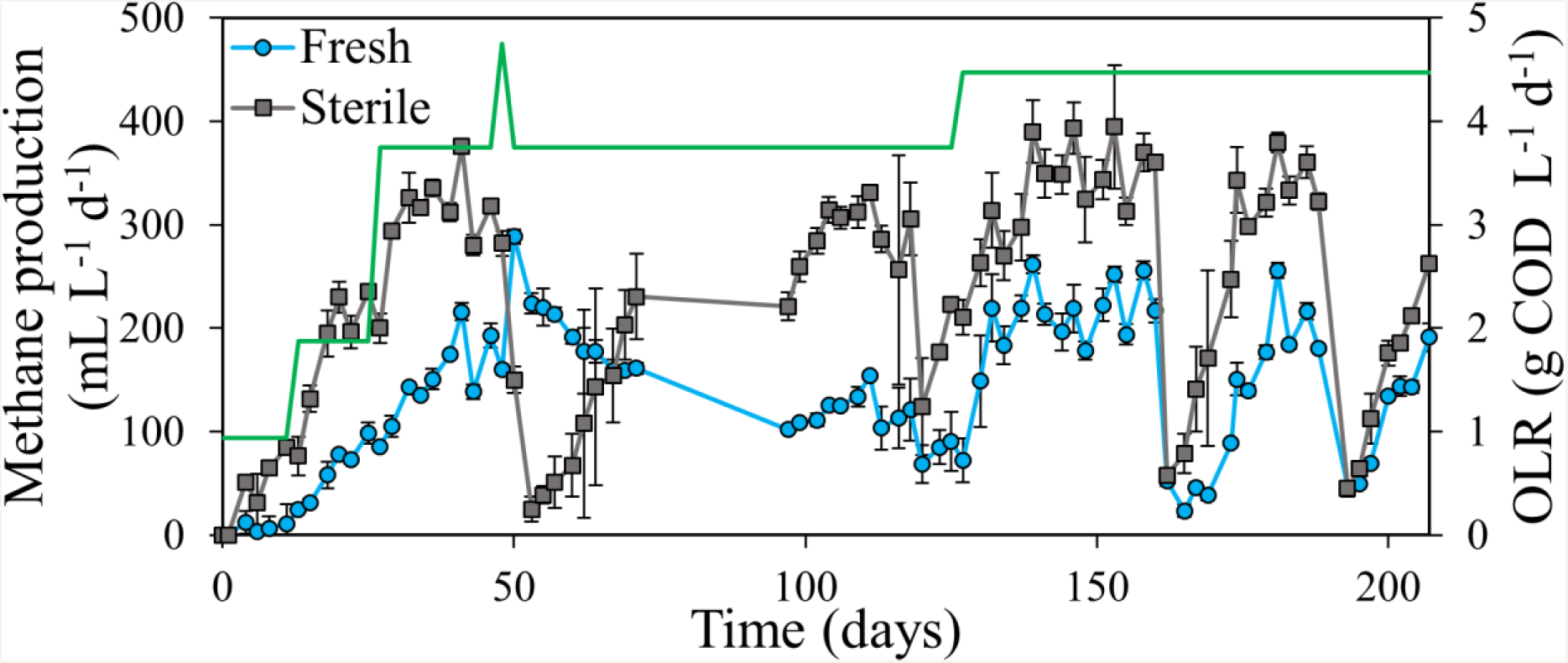
Methane production in function of time in the digesters fed fresh and sterile waste activated sludge. Average values of the biological replicates (n=3) are presented, and the error bars represent standard deviations. The green line represents the organic loading rate (OLR).

**Figure 2.**
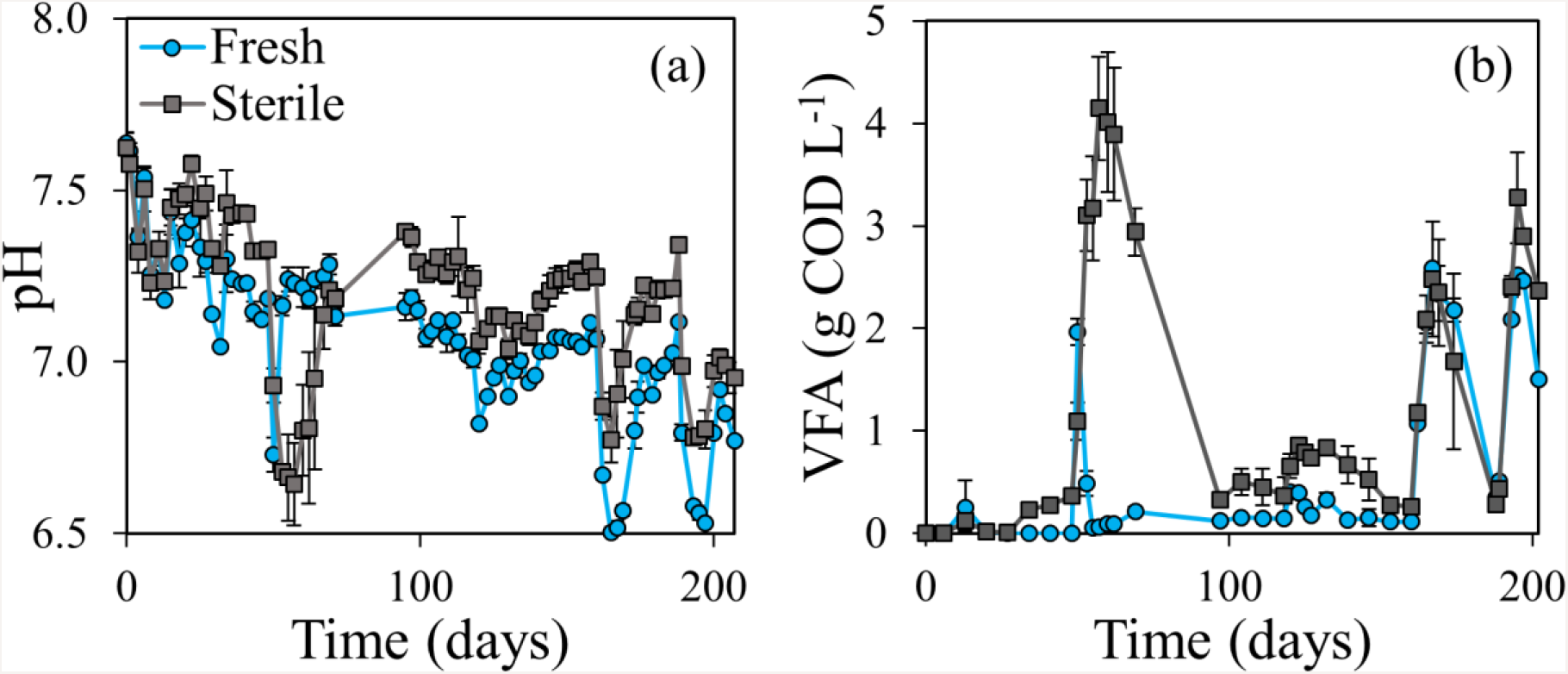
pH (a) and total volatile fatty acid (VFA) concentration (b) in function of time in the digesters fed fresh and sterile waste activated sludge. Average values of the biological replicates (n=3) are presented, and the error bars represent standard deviations.

#### 3.1.2. The impact of increasing salt pulses

After the addition of glycerol and stabilisation of the digesters to a new steady-state, NaCl was added as a novel disturbance. The increase in salt caused the increase in conductivity (Figure S2). The first pulse of 6.25 g Na^+^ L^−1^ on day 118 had only a limited, though clear influence on the methane production, pH and VFA, as a limited decrease in methane production could be observed in both digesters from day 120 on (Figure 1). A slight decrease in pH below the optimal value of 7.0 could be observed from day 120 on, but only in the digester fed fresh WAS (Figure 2a). In contrast, the residual VFA reached higher values in the digesters fed sterile WAS (Figure 2b), yet, total VFA did not exceed 1 g COD L^−1^.

A second pulse of 12.5 g Na^+^ L^−1^ on day 160 resulted in a strong inhibition of the process, as reflected in a strong decrease in methane production and pH, and an increase in residual VFA. The methane production reached similar low values of 52 ± 5 and 58 ± 6 mL CH_4_ L^−1^ d^−1^ on day 162 for the digesters fed fresh and sterile WAS, respectively (Figure 1). However, the relative decrease in methane production was higher for the digesters fed sterile WAS (84 ± 8 %) compared to the digesters fed fresh WAS (76 ± 8 %), due to the higher initial methane production in the digesters fed sterile WAS. The decrease in pH was stronger for the digesters fed fresh WAS (lowest value of 6.50 ± 0.02 on day 165) than for the digester fed sterile WAS (lowest value of 6.77 ± 0.07 on day 165), but the difference in pH between the steady-state values and the lowest value was the same for both digesters, *i.e.*, about 0.5 units (Figure 2a). The increase in residual VFA was similar for both digesters, with maximum values of 2.59 ± 0.01 and 2.48 ± 0.56 g COD L^−1^ in the digesters fed fresh and sterile WAS, respectively (Figure 2b). Complete recovery of both digesters was prior to the addition of the final pulse on day 188. The decrease in methane production and pH showed a similar trend as in response to the pulse of 12.5 g Na^+^ L^−1^ (Figure 1 & 2a). The decrease in pH was again stronger for the digesters fed fresh WAS than for the digesters fed sterile WAS (Figure 2a). The increase in residual VFA was in this case slightly higher in the digester fed sterile WAS (3.27 ± 0.44 g COD L^−1^ on day 195) than in the digester fed fresh WAS (2.52 ± 0.03 g COD L^−1^ on day 195) (Figure 2b). However, similar to the previous NaCl pulses, process recovery took place, as can be observed in the increasing biogas and pH values (Figure 1 & 2a) and decreasing residual VFA (Figure 2b) towards the end of the experiment.

### 3.2. Microbial community composition and organization

#### 3.2.1. Bacterial community

An average of 13,774 ± 2,770 reads across all samples, representing 1,488 ± 520 OTUs were obtained per sample (including singletons) following amplicon sequencing. Removal of singletons and normalisation according to the common-scale approach resulted in an average of 9,182 ± 327 reads and 648 ± 240 OTUs per sample. No significant differences (repeated measures ANOVA, *P* < 0.0001) could be detected in the bacterial community composition between the biological replicates.

The bacterial community composition strongly differed between the digesters fed fresh and sterile WAS (PERMANOVA, *P* = 0.0001), with 941 OTUs (21.7 % of all OTUs, reflecting 86.6 ± 6.1 % of the total relative abundance) showing a significant difference (*DESeqDataSetFromMatrix*, *P* < 0.05) in relative abundance, irrespective of the salt concentration or time point. This difference was already clearly visible in the four main phyla, with the *Actinobacteria* (14.7 ± 2.0 vs. 4.7 ± 5.3 %) and *Proteobacteria* (30.5 ± 1.0 vs. 2.6 ± 2.5 %) showing a higher relative abundance in the digesters fed fresh WAS, and the *Bacteroidetes* (25.9 ± 5.6 vs. 16.3 ± 2.7 %) and *Firmicutes* (44.0 ± 7.1 vs. 14.6 ± 4.5 %) showing a higher relative abundance in the digesters fed sterile WAS (Figure 3). This difference in bacterial community composition was also clear on the class, order, family, and OTU level (Figure S3-S6). The α-diversity analysis revealed a significantly higher richness H_0_ (*P* = 0.0025), and overall diversity H_1_ (*P* = 0.0015) and H_2_ (*P* = 0.0015) in the digesters fed fresh WAS, compared to the digesters fed sterile WAS (Figure 4).

**Figure 3.**
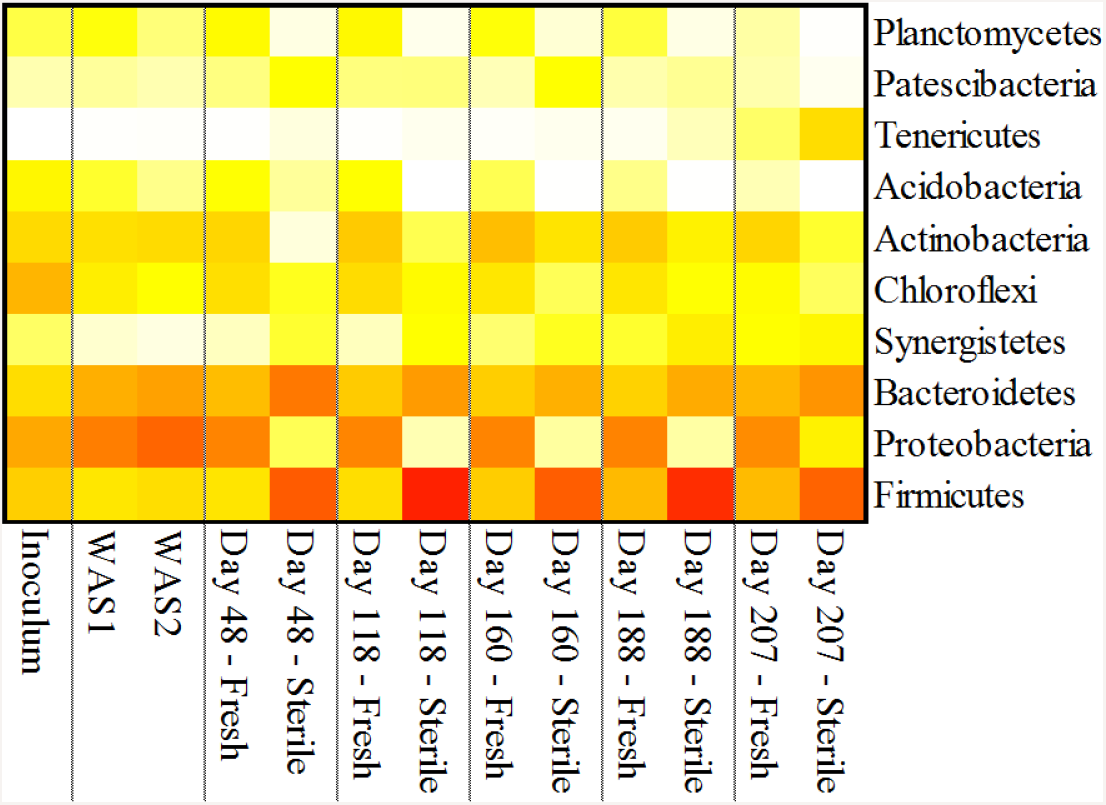
Heatmap showing the relative abundance of the bacterial community at the phylum level in the inoculum, two batches waste activated sludge (WAS1 & 2) and in the digesters fed fresh and sterile waste activated sludge on day 48, 118, 160, 188 and 207. Weighted average values of the biological replicates (n=3) are presented. Only those phyla with at least 1 % relative in one of the samples are included. The colour scale ranges from 0 (white) to 60% (red) relative abundance.

**Figure 4.**
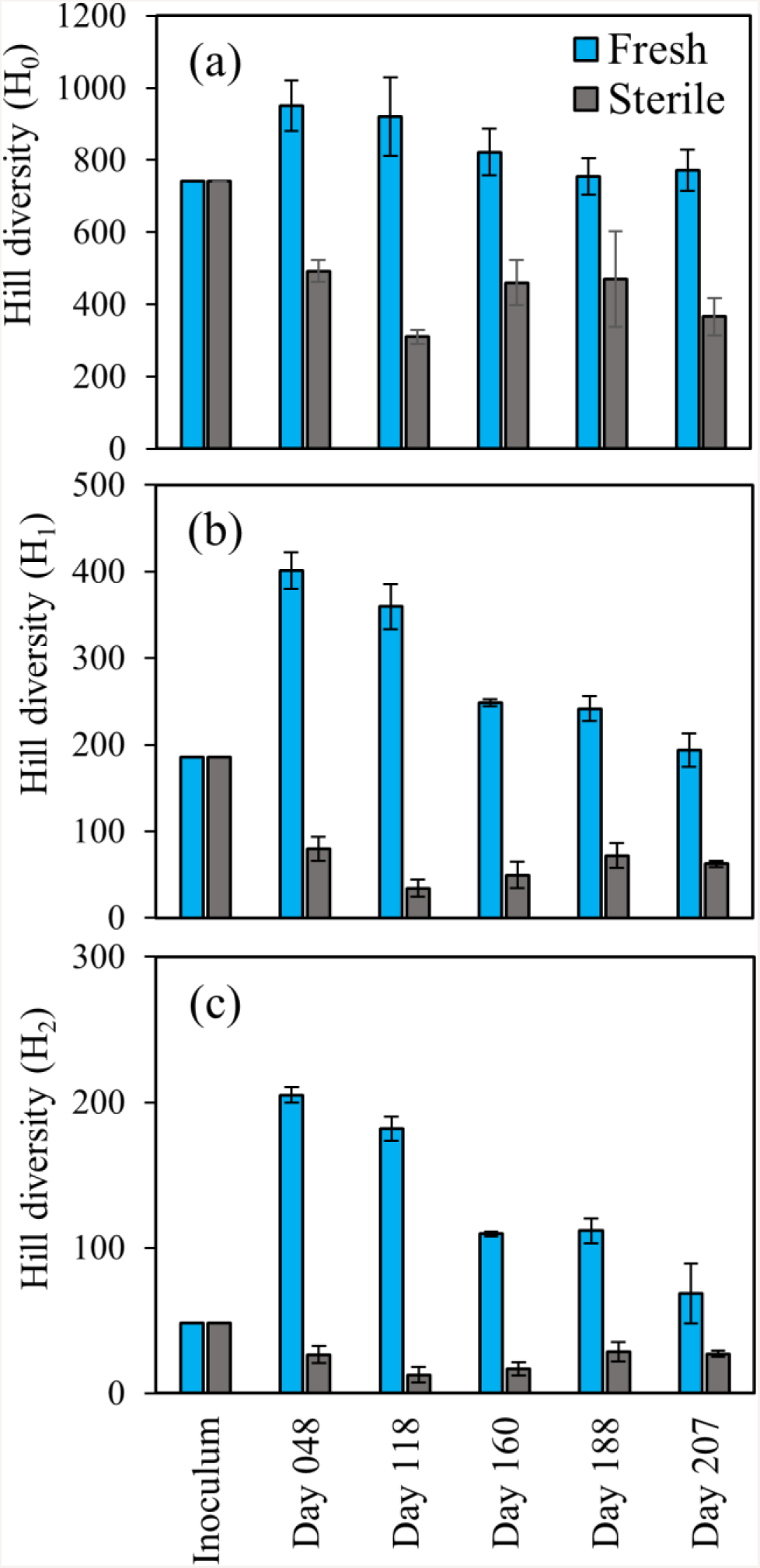
Alpha diversity of the inoculum and in the digesters fed fresh and sterile waste activated sludge on day 48, 118, 160, 188 and 207. The three Hill order diversity numbers (a) H0 (richness, number of OTUs), (b) H1 (exponential value of the Shannon index) and (c) H2 (inverse Simpson index) were calculated based at the OTU level. Error bars represent standard deviations of the biological replicates (n=3).

Even though the addition of glycerol and sodium impacted the overall methane production process, its direct effect on the bacterial community was limited. In total, 336 OTUs (7.7 % of all OTUs) showed a significant (*DESeqDataSetFromMatrix*, P < 0.05) increase or decrease in function of the increasing sodium doses. Although the shift in relative abundance in response to the increased salinity could be detected for several dominant OTUs, such as OTU00004 (unclassified *Rikenellaceae*) and OTU00007 (unclassified Bacterium) (Figure S6), this shift was not observed on the different phylogenetic levels (Figure 3 & S3-S5). A differential impact of the increased sodium concentration on α-diversity could be observed between the digesters fed fresh and sterile WAS (Figure 4). For the digesters fed fresh WAS, a significant negative correlation was observed between the sodium concentration and H0 (τ = −0.47, *P* = 0.021), H1 (τ = −0.69, *P* = 0.0006) and H2 (τ = −0.67, *P* = 0.0010) diversity. In contrast, the increasing sodium concentration did not seem to impact the H0 ((τ = −0,06 *P* = 0.77), H1 (τ = 0.11, *P* = 0.55) and H2 (τ = 0.31, *P* = 0.11) diversity in the digesters fed sterile WAS.

#### 3.2.2. Methanogenic community

Real-time PCR analysis of the total bacteria and different methanogenic populations revealed a similar methanogens:bacteria ratio of 0.42 ± 0.23 % across all digester samples, excluding the WAS samples. This indicates an overall strong dominance of the bacteria over the methanogens (at least a factor 100 higher absolute abundance) in the microbial community. No clear effect could be observed related to the feedstock sterilisation, although the methanogens:bacteria ratio appeared to be higher in the digesters fed sterile WAS, especially in response to the salt pulses (Figure S7). The methanogenic community in the two WAS batches was similar, with a dominance of the *Methanosaetaceae*, and this was also the case for the Inoculum (Figure 5 and S8).

**Figure 5.**
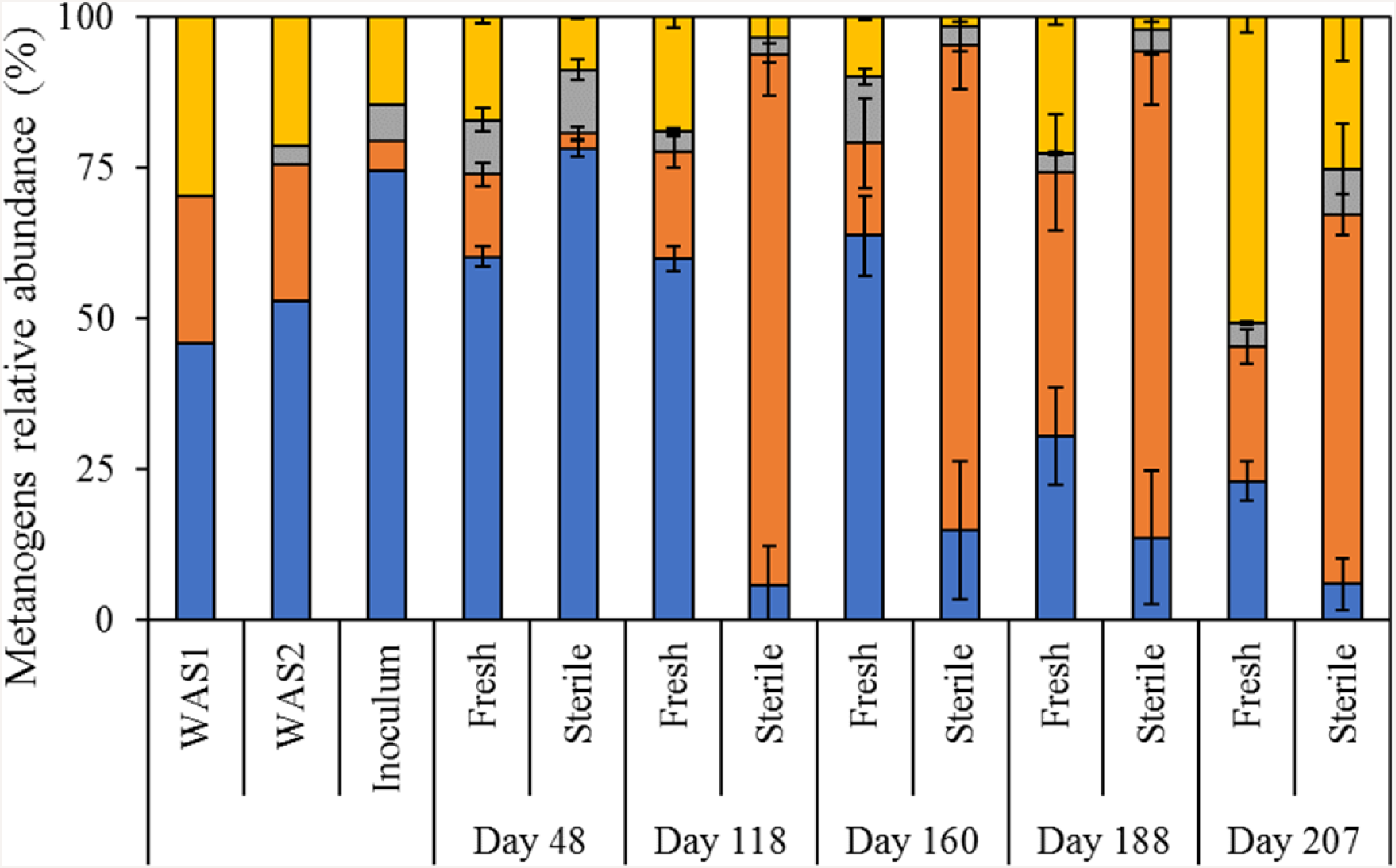
Relative abundance (%) of the *Methanosaetaceae* (blue, 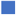), *Methanosarcinaceae* (orange, 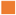), *Methanobacteriales* (grey, 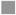) and *Methanomicrobiales* (yellow, 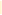) in the methanogenic community of the two batches waste activated sludge (WAS1 & 2), the inoculum and in the digesters fed fresh and sterile waste activated sludge on day 48, 118, 160, 188 and 207. Average values of the biological replicates (n=3) are presented, and the error bars represent standard deviations.

Overall, a significant difference (PERMANOVA, *P* = 0.0003) in the methanogenic community profile could be observed between the digesters fed fresh and sterile WAS. After 48 days, prior to the addition of glycerol or sodium, a first divergence between the digesters with fresh and sterile WAS could be observed, with an increase in relative and absolute abundance of the *Methanosarcinaceae* in the digesters fed fresh WAS (13.6 ± 1.9 %), compared with the digesters fed sterile WAS (2.6 ± 1.0 %) (Figure 5). The first and only pulse of glycerol, prior to the addition of sodium (day 118), did not provoke a clear effect on the methanogenic community in the digesters fed fresh WAS. In contrast, the *Methanosarcinaceae* showed a clear increase in relative abundance (88.0 ± 6.7 %) on day 118 in the digesters fed sterile WAS, which was mainly due to the decrease in absolute abundance of the other methanogenic populations (Figure S8). This increase in relative and absolute abundance of the *Methanosarcinaceae* was maintained in the digesters fed sterile WAS in response to the increasing sodium pulses (day160, 188 and 207). In contrast, even though the *Methanosarcinaceae* also increased in relative and absolute abundance in the digesters fed fresh WAS in response to the increasing sodium pulses, the *Methanomicrobiales* became dominant on day 188 and especially day 207, reaching a relative abundance of 50.7 ± 2.6 %. The *Methanomicrobiales* became also more dominant in the digesters fed sterile WAS, though the *Methanosarcinaceae* remained their dominancy.

## 4. Discussion

Inoculation of the microbial community in anaerobic digestion *via* the feedstock, in this case waste activated sludge, resulted in a differentiating impact with respect to resistance to disturbances. The microbial community in the feedstock positively contributed resistance to organic overloading, in contrast to applying a sterile feedstock. However, a collapse in methane production in response to increasing salt pulses could not be prevented through feedstock inoculation, and process recovery took place irrespective of an active microbial community in the feedstock. Feedstock sterilisation strongly impacted the microbial community in the digesters in terms of composition and organisation.

### 4.1. Different disturbances have a differentiating effect on process stability in relation to feedstock inoculation

The addition of glycerol and salt showed a different impact on the activity of the microbial community. Glycerol that was added as an extra carbon source (Ma et al. 2008) can have two different effects on the process, which depends on the concentration, *i.e.*, (1) an increase in methane production or (2) process failure due to overloading *(Fountoulakis et al. 2010; González Arias et al. 2018; Holm-Nielsen et al. 2008)*. The addition of salt will normally inhibit microbial activity (Appels et al. 2008), but in this research only a temporal effect on microbial activity and process performance could be observed. When glycerol was added into the digesters, the recovery of the digester fed fresh WAS was faster than for the digesters fed sterile WAS. The reason behind this difference could be related to the presence of an active microbial community in the WAS (Lebiocka et al. 2018; Li et al. 2018), while this was not the case for the digesters fed sterile WAS. The presence of the active community in the WAS feedstock contributed to the degradation of the organics, which led to an elevated methane production (Fountoulakis et al. 2010; González Arias et al. 2018). Instead of an elevated methane production, a decrease was observed in the digesters fed sterile WAS, caused by the incremental organic loading because of glycerol addition. The microorganisms present in the digesters will quickly convert these organics into VFA, whereby the degradation of glycerol into VFA, mainly propionate and acetate, is faster than their subsequent conversion to methane. Hence, the methanogens could not keep up with the conversion rate of glycerol to VFA, which resulted in the accumulation of VFA (González Arias et al. 2018; Holm-Nielsen et al. 2008). This accumulation was confirmed by the pH decrease. Not only the addition of glycerol caused the accumulation of VFA, but also the sterilisation of the feed had an impact on the formation of VFA. The sterilisation of the influent resulted in an increase in the soluble COD fraction, as also reported earlier (Papadimitriou 2010), mainly due to the increase in VFA. The COD that normally was not biodegradable in the influent was partially converted into readily biodegradable COD, mainly VFA, because of the sterilisation process (Papadimitriou 2010; Tampio et al. 2014). Combined with the addition of glycerol, this resulted in an elevated organic loading, which led to the accumulation of VFA, and eventually to the inhibition of the microorganisms (Shi et al. 2017). Hence, the differential effect can be attributed to the presence of microbial community with a higher degree of activity in the fresh WAS and/or the increase in readily biodegradable COD in the sterile WAS.

### 4.2. Process recovery takes place irrespective of increasing salt pulses

With the addition of salt, no extra COD was introduced into the digesters, but salt is known as a common inhibitor of microbial activity in AD (Chen et al. 2008; Rinzema et al. 1988; Zhang et al. 2017a). The addition of salt commonly has a stronger impact, by causing a higher osmotic imbalance, on the methanogenic archaea, in comparison to the bacterial community (Rinzema et al. 1988; Wang et al. 2017; Zhang et al. 2017b). Despite elevated concentrations of NaCl, the bacterial community remained active, whereby they still converted the organics into VFA. In contrast, the salt concentration, especially at 12.5 and 25.0 g Na^+^ L^−1^, temporarily ceased methanogenic activity, resulting into an accumulation of residual VFA (Appels et al. 2008; Chen et al. 2008). The inhibition was only temporarily, because the methanogens managed to recover following the decrease in salt (Feijoo et al. 1995; Hierholtzer and Akunna 2014; Ismail et al. 2010), which is reflected in the decrease of VFA, and increases in pH and methane production. Because of the rather low organic loading rate and limited biodegradability of the WAS, the VFA accumulation potential was low, and residual VFA increase and pH decrease remained limited in both cases. Therefore, based on the results of both disturbances, *i.e.*, glycerol and NaCl, it could be hypothesized that long-term inhibition in AD is not caused by the cations themselves, which only temporarily inhibit methanogenesis, but rather by the subsequent accumulation of VFA that permanently inhibit methanogenesis at higher organic loading rates (Mischopoulou et al. 2017; Zhang et al. 2017a). The similarity of both treatments demonstrates that the microorganisms in the feedstock apparently did not have an influence on the inhibition caused by salts. This indicates that process inhibition in AD is a consequence of a tilting balance in terms of VFA accumulation, as a consequence of another disturbance, resulting in a pH drop (Kugelman and McCarty 1965), rather than being directly caused by the disturbance. This hypothesis remains to be confirmed through other disturbances.

### 4.3. Feedstock inoculation determines microbial community composition and organisation

The microbial community in AD, containing representatives from both the bacterial and archaeal domains of life, is determined by different parameters central to the performance of the process, including total and free ammonia, temperature, salinity and pH (De Vrieze et al. 2018; De Vrieze et al. 2015b; Garcia and Angenent 2009; Westerholm and Schnürer 2019). This was also reflected in our study, as especially the addition of multiple salt pulses affected the bacterial and archaeal microbial community. However, this effect was clearly overshadowed by the differential impact of using fresh vs. sterile WAS as feedstock. This observation emphasises the importance of the feedstock composition with respect to the microbial community on two levels. First, even though certain operational parameters, such as temperature and solids retention time, can be set by the operator, the feedstock composition determines the microbiome (Sundberg et al. 2013; Zhang et al. 2014), related to the abovementioned parameters. In our study, the impact of sterilisation of the feed, which also changed the WAS chemical composition, on the process itself at stable conditions was obvious, related to the increase in methane production, pH and residual VFA at steady-state conditions. Second, the influx of microorganisms can be a key factor that influences or even steers the AD process (Kirkegaard et al. 2017; Shin et al. 2019), as observed in our study in response to a glycerol pulse. Sterilisation through autoclaving, in addition to killing of all microorganisms, also should degrade DNA molecules into small fragments (20-30 base pairs), though some larger fragments can remain behind (Esser et al. 2006), especially related to the complex WAS matrix. The WAS contains a living active microbial community with both bacteria and methanogenic archaea, as demonstrated earlier (De Vrieze et al. 2015a), but also dead/inactive microorganisms and/or free DNA could be present, thus, influencing the microbial community profile (Kirkegaard et al. 2017). However, given (1) the presence of anaerobic sites in activated sludge units, reflected in diffuse methane emissions (Daelman et al. 2012), and (2) the importance of immigration in microbial community shaping (Sloan et al. 2006), the contribution of microorganisms in the WAS feedstock to process performance in AD is apparent, as confirmed by our results.

The impact of feedstock inoculation on the microbial community can be considered on different levels and across the domains of life. The strong difference in bacterial community composition already at the phylum level can be partially contributed directly to the WAS. For example, the digesters fed fresh WAS contained a higher relative abundance of Proteobacteria, thus, reflecting the dominance of this phylum in the WAS itself. This was also reflected on the other phylogenetic levels. A similar result was obtained for the methanogenic community, because even though the absolute abundance of the methanogens in the WAS was up to a factor 10-100 lower than in the digesters, the digesters fed fresh WAS showed a similar profile as the WAS itself during the first sampling points. The strong dominance of the *Methanosarcinaceae* in the digesters fed sterile WAS could be due to their high growth rate at higher residual VFA (Conklin et al. 2006) as r-strategist in a more “open” niche (De Vrieze et al. 2017; Pianka 1970) in contrast to the digesters fed fresh WAS, where there was a continuous inflow of methanogens. The dominance of fewer taxa in the more open niche of the digesters fed sterile WAS was also reflected in their significantly lower diversity, compared to the digesters fed fresh WAS. As higher overall diversity can be linked to a higher functional redundancy (Briones and Raskin 2003; Langer et al. 2015; McMahon et al. 2007; Venkiteshwaran et al. 2016), this, at least partially, explains the higher resistance to the glycerol pulse of the digesters fed fresh WAS, compared to the digesters fed sterile WAS.

## 5. Conclusions

We demonstrated the importance of feedstock inoculation with respect to process performance and resistance towards disturbances in anaerobic digestion. A differential effect of feedstock inoculation was observed, as the microorganisms in the feedstock contributed to process stability in response to a glycerol pulse, while this was not the case for increasing salt pulses. Process recovery following the salt pulses took place irrespective of inoculation *via* the feedstock. Feedstock inoculation strongly determined the bacterial and archaeal community composition and organisation in the digesters, providing additional process stability security. Overall, this opens the need to consider feedstock inoculation with feedstocks rich in suitable microorganisms for anaerobic digestion, such as waste activated sludge and manure, for full-scale anaerobic digestion processes as a “right-of-the-shelve” strategy to enhance process stability.

## Supporting information

Supplementary file 1 containing all additional Materials and Methods and Results

Supplementary file 2 Raw OTU Table

## Acknowledgments

Cindy Ka Y Law kindly acknowledges the support from the ELECTRA project “Electricity Driven Low Energy and Chemical Input Technology for Accelerated Bioremediation” financed by the H2020 of the European Commission under Grant number GA 826244. Jo De Vrieze is supported as postdoctoral fellow by the Research Foundation Flanders (FWO-Vlaanderen). The authors would like to thank Tim Lacoere for his contribution to the molecular analyses and Aquafin for their assistance with sample collection. We thank Pieter Candry and Karel Folens for critically reading the manuscript.

