## Supplementary file 1 containing all additional Materials and Methods and Results for "The feedstock microbiome selectively steers process stability during the anaerobic digestion of waste activated sludge"

Journal: Applied Microbiology and Biotechnology

### S1. Characterisation fresh vs. sterile waste activated sludge

**Table S1** Comparison of the two batches of waste activated sludge (n=3) before and after sterilisation trough autoclavation (twice) for 30 min at 121 °C. TS = total solids, VS = volatile solids, COD = chemical oxygen demand, VFA = volatile fatty acids, TAN = total ammonia nitrogen, TKN = Kjeldahl nitrogen, FW = fresh weight.

| | | Fresh WAS | | Sterile WAS | |
|---|---|---|---|---|---|
| Parameter | Unit | WAS1 | WAS2 | WAS1 | WAS2 |
| pH | - | 7.04 ± 0.01 | 6.37 ± 0.02 | 8.00 ± 0.01 | 7.63 ± 0.35 |
| TS | g TS $kg^{-1}$ FW | 76.2 ± 0.9 | 54.3 ± 0.1 | 67.7 ± 1.9 | 48.1 ± 0.3 |
| VS | g VS $kg^{-1}$ FW | 44.2 ± 0.7 | 33.6 ± 0.1 | 31.7 ± 1.1 | 19.2 ± 0.1 |
| Soluble COD | g COD $kg^{-1}$ FW | 0.8 ± 0.2 | 2.3 ± 1.1 | 5.9 ± 0.3 | 7.2 ± 0.8 |
| Conductivity | mS $cm^{-1}$ | 4.69 ± 0.01 | 4.30 ± 0.01 | 7.00 ± 0.10 | 7.58 ± 0.04 |
| Total VFA | g COD $kg^{-1}$ FW | 0 ± 0 | 0 ± 0 | 7.0 ± 0.7 | 12.6 ± 0.3 |
| TAN | mg N $kg^{-1}$ FW | 345 ± 13 | 571 ± 7 | 967 ± 27 | 1045 ± 32 |
| $Na^+$ | mg $Na^+$ $kg^{-1}$ | 173 ± 44 | 67 ± 17 | 109 ± 2 | 62 ± 8 |
| $K^+$ | mg $K^+$ $kg^{-1}$ | 164 ± 1 | 183 ± 3 | 181 ± 4 | 200 ± 27 |
| TS:VS ratio | - | 1.72 ± 0.03 | 1.62 ± 0.01 | 2.13 ± 0.10 | 2.50 ± 0.02 |

### S2. Amplicon sequencing and data processing

S2.1. Amplicon sequencing

The primers 341F (5’- CCTACGGGNGGCWGCAG) and 785R (5’-GACTACHVGGGTATCTAATCC) that target the V3-V4 region of the 16S rRNA gene (Klindworth et al. 2013), with an extra wobble position in the reverse primer to make it more universal, were used to target total bacteria. The DNA extracts had a minimum DNA concentration of 10 ng $\mu L^{-1}$, and were free of RNA, which was validated with agarose gel electrophoresis. A 2-step PCR protocol was used, and 10-25 ng of DNA was used as template for the first PCR run with a total volume of 50 µL, using the 341F and the 785R primers with Illumina adaptor sequences. The first PCR run contained 25 cycles, with an annealing temperature of 55 °C. The PCR products were purified using Ampure XP beads according to the manufacturer’s instructions, and their size checked on a Fragment analyser (Advanced Analytical Technologies, Inc., Ames, Iowa, USA), and quantified by fluorometric analysis. The purified PCR products were subjected to a 2$^{nd}$ PCR run that contained 6 cycles with an annealing temperature of 55 °C, using sample-specific barcoded primers (Nextera XT index kit, Illumina). The PCR products were again purified using Ampure XP beads according to the manufacturer’s instructions, and their size checked on a Fragment analyser, and quantified. Next, multiplexing, clustering and sequencing was carried out on an Illumina MiSeq with the paired-end (2×) 300 bp protocol and indexing. The sequencing run was analysed *via* the Illumina CASAVA pipeline (v1.8.3) in which demultiplexing was based on sample-specific barcodes. The raw sequencing data were processed, removing sequence reads of too low quality (only "passing filter" reads were selected), and discarding reads containing adaptor sequences or PhiX control with an in-house filtering protocol. A quality assessment on the remaining reads was performed using the FASTQC quality control tool version 0.10.0.

S2.2. Data processing

The Mothur software package (v.1.40.3), and guidelines developed by Schloss et al. (2009) were used to process the raw Illumina data on a GNU/Linux 3.16.0-46-generic x86_64 system. The forward and reverse reads were assembled into contigs by a heuristic approach, taking the Phred quality scores into account. Ambiguous contigs or with unsatisfying overlap were removed, and the remaining sequences were aligned to the mothur formatted silva seed v132 database. Those sequences not aligning within the region targeted by the primer set or sequences with homopolymer stretches with a length > 12 bp were removed. The sequences were pre-clustered, allowing mismatch for every 100 bp of sequence. Chimeric sequences were removed with UCHIME (Edgar et al. 2011). Classification of the sequences was carried out by a naïve Bayesian classifier, using the RDP 16S rRNA gene training set, release 16, with an 85% cut-off for the pseudobootstrap confidence score. Taxa that were annotated as Chloroplast, Mitochondria, unknown, Archaea or Eukarya at the kingdom level were excluded. Sequences were clustered into OTUs with an average linkage, and at a 97% sequence identity, using the OptiClust method (Westcott and Schloss 2017). Representative sequences were picked for each OTU as the most abundant sequence within that OTU.

### S3. Real-time PCR analysis

Real-time PCR (qPCR) analysis was carried out on a StepOne-Plus$^{TM}$ Real-Time PCR System (Applied Biosystems, Carlsbad, CA). The DNA extracts were diluted with PCR water (DNase free) to a final template DNA concentration between 1-10 ng µL$^{-1}$, and real-time PCR analysis was carried out in technical triplicates. The different methanogenic orders *Methanobacteriales* (MBT857F and MBT1196R) and *Methanomicrobiales* (MMB282F and MMB832R), and the families *Methanosaetaceae* (Mst702F and Mst862R) and *Methanosarcinaceae* (Msc380F and Msc828R) were quantified using the primer sets and protocol described by Yu et al. (2005). Total bacteria were quantified using the P338f and P518r primers (Ovreas et al. 1997).

The PCR reaction mixture (20 µL) was prepared with the iTaq$^{TM}$ Universal SYBR$^{®}$ Green Supermix (Bio-Rad Laboratories, Hercules, CA, USA), and contained 10 µL of 2× iTaq$^{TM}$ universal SYBR$^{®}$ Green supermix, 0.8 µL of the forward and reverse primer (10 µM stock solution), 6.4 µL of nuclease-free water and 2 µL of template DNA.

The PCR program was carried out in a two-step thermal cycling process. For the methanogenic orders *Methanobacteriales* and the families *Methanosaetaceae* and *Methanosarcinaceae*, the program consisted of a predenaturation step of 10 min at 94 °C, followed by 40 cycles of 10 s at 94 °C and 1 min at 60 °C. The annealing temperature was set at 63 °C for the *Methanomicrobiales* order. To quantify total bacteria, the denaturation step lasted 15 s. Standard curves were generated using the DNA extracts of the pure cultures *Methanosaeta concilli* (DSM2139) for the *Methanosaetaceae*, *Methanosarcina barkeri* (DSM 800) for the *Methanosarcinaceae*, *Methanobrevibacter arboriphilicus* (DSM 1536) for the *Methanobacteriales*, and *Methanomicrobium mobile* (DSM 1539) for *Methanomicrobiales*. Linearized plasmids were used to create a tenfold dilution series of $10^1$–$10^7$ copies µL$^{-1}$, and these were analysed in triplicate. The real-time PCR data were represented as copies per gram of wet sludge. Real-time PCR quality was evaluated by means of the different parameters obtained through analysis with the StepOnePlus software V2.3 (Table S2).

**Table S2** Quality control of the parameters for real-time PCR analysis. These parameters were obtained during analysis with the StepOnePlus V2.3 software. The detection limit was calculated as copies of the target 16S rRNA gene fragment per gram wet sludge, and was determined taking both dilution and extraction efficiency into account. The represented values are average values and standard deviations of the two PCR plates that were used to analyse all samples.

| Parameter | Slope | $R^2$ | Efficiency (%) | Detection limit (copies $g^{-1}$) |
|---|---|---|---|---|
| *Methanosaetaceae* | -3.6 | 1.00 | 88.0 | $2.52 \times 10^4$ |
| *Methanosarcinaceae* | -3.7 | 1.00 | 87.8 | $2.37 \times 10^4$ |
| *Methanobacteriales* | -4.1 | 1.00 | 75.9 | $2.20 \times 10^4$ |
| *Methanomicrobiales* | -3.9 | 1.00 | 80.5 | $1.88 \times 10^4$ |
| Total Bacteria | -3.6 | 1.00 | 89.4 | $1.05 \times 10^5$ |

### S4. Rarefaction curves

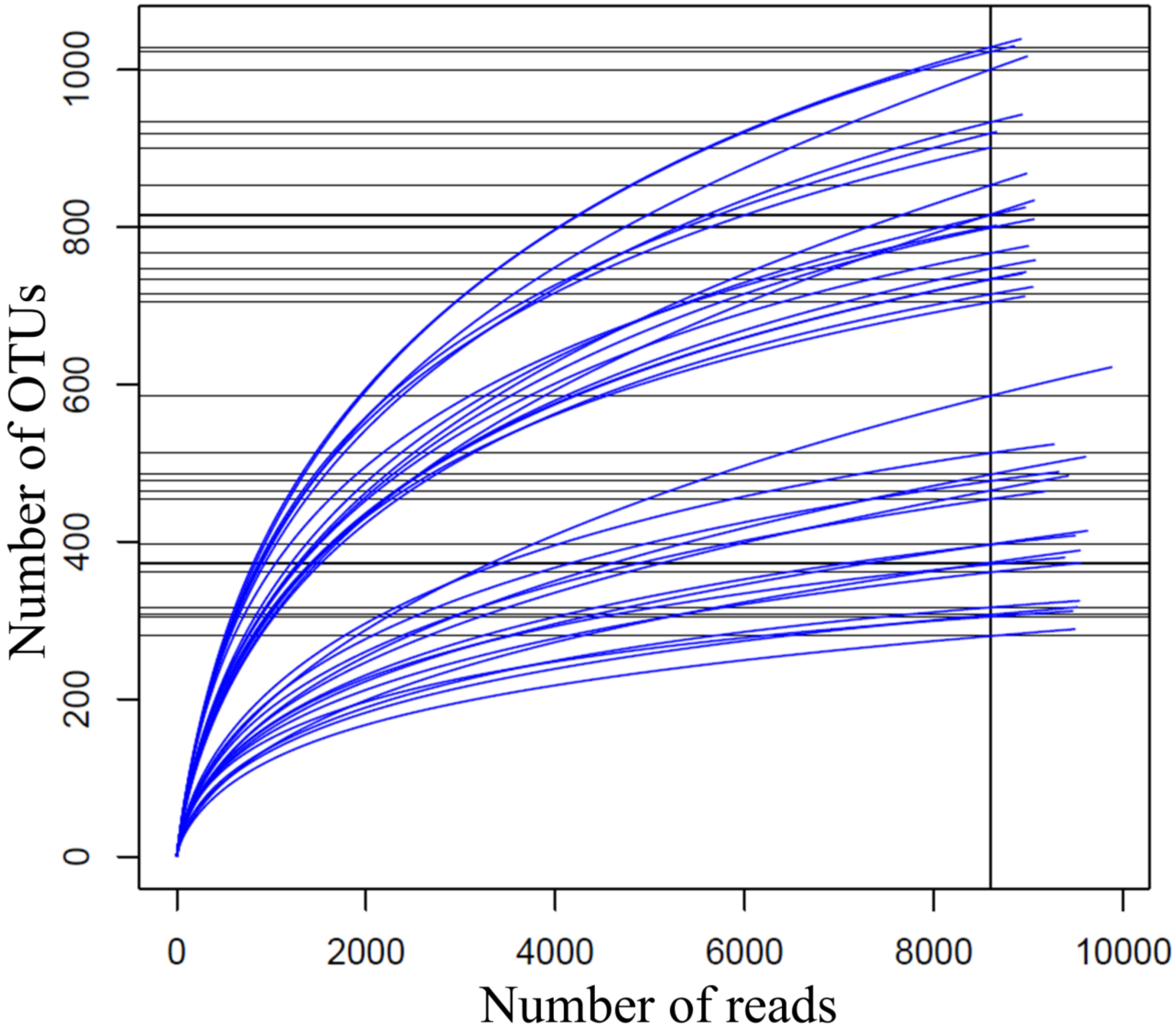

**Figure S1** Rarefaction curves indicating the number of resolved OTUs against sampling depth of each of the samples after removal of singletons and normalisation according to the common-scale approach.

### S5. Biogas composition

The composition of the biogas was determined by means of a Compact GC (Global Analyser Solutions, Breda, The Netherlands). The first channel, using He as carrier gas, consisted of a double channel with a Porabond Q precolumn and Molsieve 5A column for $CH_4$ analysis and a Rt-QSBond precolumn and Rt-QSBond column for $CO_2$ and $H_2S$ measurement. The second channel, using $N_2$ as carrier gas, with a Porabond Q precolumn and Molsieve 5A column was used to measure $H_2$. Both channels had a sample loop of 25 µL. The thermal conductivity detector had a lower detection limit of 100 ppmv for each gas component.

### S6. Volatile fatty acid analysis

The VFA concentrations (C2-C8) were measured with a gas chromatograph (GC-2014, Shimadzu®, The Netherlands) that was equipped with a DB-FFAP 123-3232 column (30 m x 0.32 mm x 0.25 µm; Agilent, Belgium) and a flame ionization detector (FID). The samples (2 mL) were conditioned with sulphuric acid and sodium chloride, and 2-methyl hexanoic acid was used as internal standard to quantify the extraction with diethyl ether. The extracted sample (1 µL) was injected at 200 ºC with a split ratio of 60 and a purge flow of 3 mL $min^{-1}$. The oven temperature increased with 6 ºC $min^{-1}$ from 110 ºC to 165 ºC, where it was kept for 2 min. The FID had a temperature of 220 ºC. The carrier gas was $N_2$ at a flow rate of 2.49 mL $min^{-1}$. The detection limit was 30 mg $L^{-1}$ for acetate, 10 mg $L^{-1}$ for propionate and 2 mg $L^{-1}$ for the other VFA (C4-C8).

### S7. Operational parameters

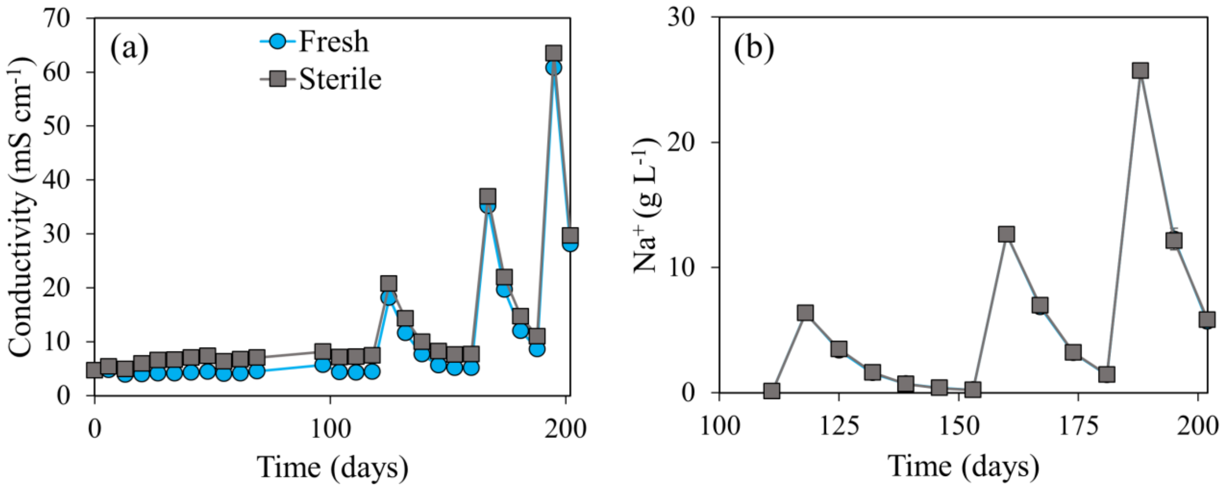

**Figure S**2 The conductivity (a) and $Na^+$ concentration (b) in the digesters fed fresh and sterile waste activated sludge. Average values of the biological replicates (n=3) are presented, and the error bars represent standard deviations.

### S8. Heatmaps

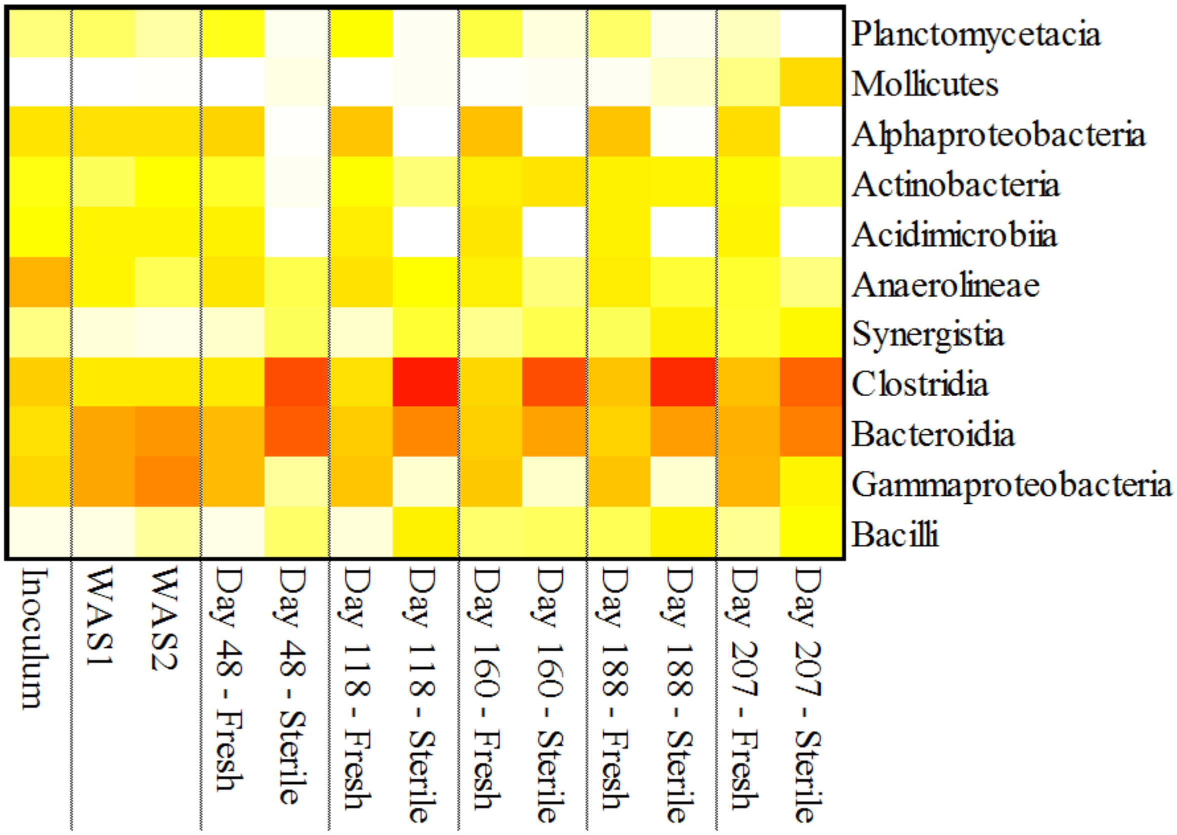

**Figure S3** Heatmap showing the relative abundance of the bacterial community at the class level in the inoculum, two batches waste activated sludge (WAS1 & 2) and in the digesters fed fresh and sterile waste activated sludge on day 48, 118, 160, 188 and 207. Weighted average values of the biological replicates (n=3) are presented. Only those classes with at least 1 % relative abundance in one of the samples are included. The colour scale ranges from 0 (white) to 50% (red) relative abundance.

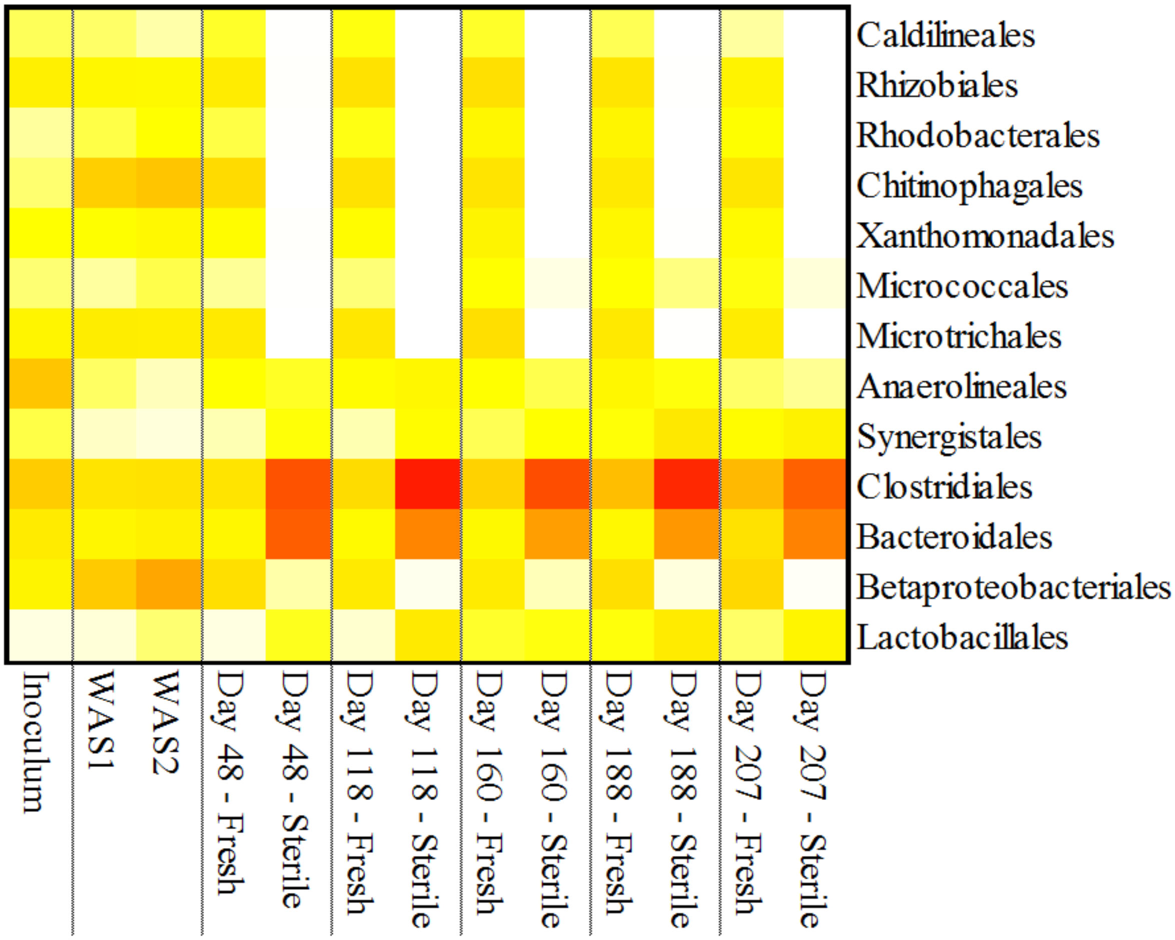

**Figure S4** Heatmap showing the relative abundance of the bacterial community at the order level in the inoculum, two batches waste activated sludge (WAS1 & 2) and in the digesters fed fresh and sterile waste activated sludge on day 48, 118, 160, 188 and 207. Weighted average values of the biological replicates (n=3) are presented. Only those orders with at least 1 % relative abundance in one of the samples are included. The colour scale ranges from 0 (white) to 50% (red) relative abundance.

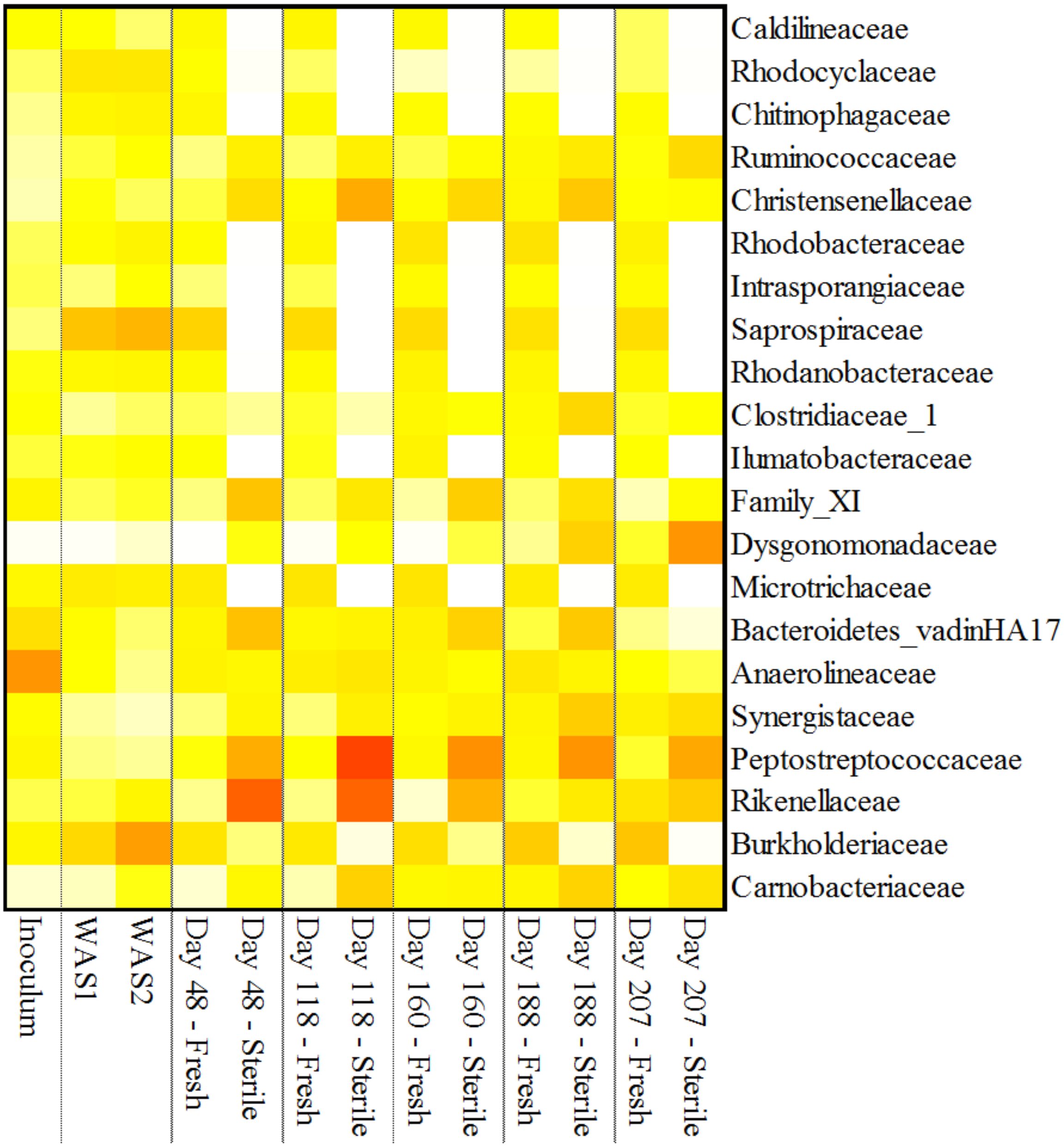

**Figure S5** Heatmap showing the relative abundance of the bacterial community at the family level in the inoculum, two batches waste activated sludge (WAS1 & 2) and in the digesters fed fresh and sterile waste activated sludge on day 48, 118, 160, 188 and 207. Weighted average values of the biological replicates (n=3) are presented. Only those families with at least 1 % relative abundance in one of the samples are included. The colour scale ranges from 0 (white) to 30% (red) relative abundance.

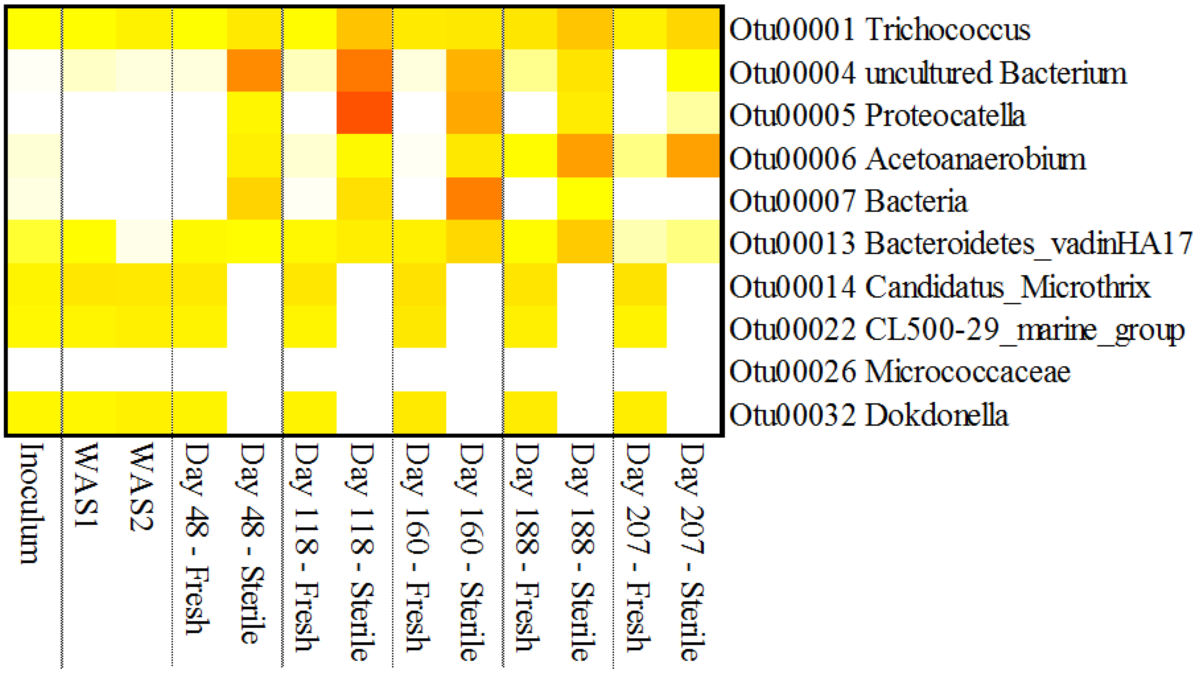

**Figure S6** Heatmap showing the relative abundance of the bacterial community at the OTU level in the inoculum, two batches waste activated sludge (WAS1 & 2) and in the digesters fed fresh and sterile waste activated sludge on day 48, 118, 160, 188 and 207. Weighted average values of the biological replicates (n=3) are presented. Only those OTUs with at least 1 % relative abundance in one of the samples are included. The colour scale ranges from 0 (white) to 30% (red) relative abundance.

### S9. Real-time PCR results

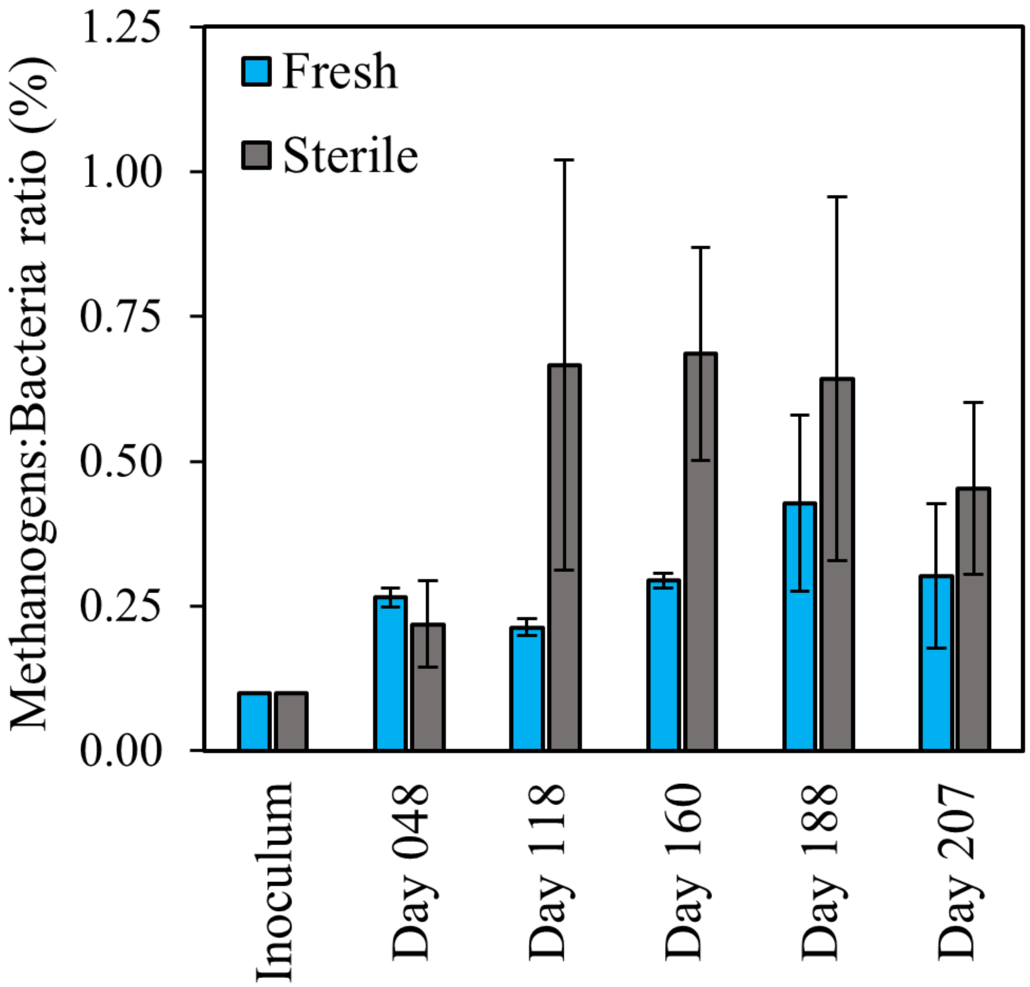

**Figure S7** Ratio total methanogens to total bacteria in the inoculum and the digesters fed fresh and sterile waste activated sludge on day 48, 118, 160, 188 and 207. Error bars represent standard deviations of the biological replicates (n=3).

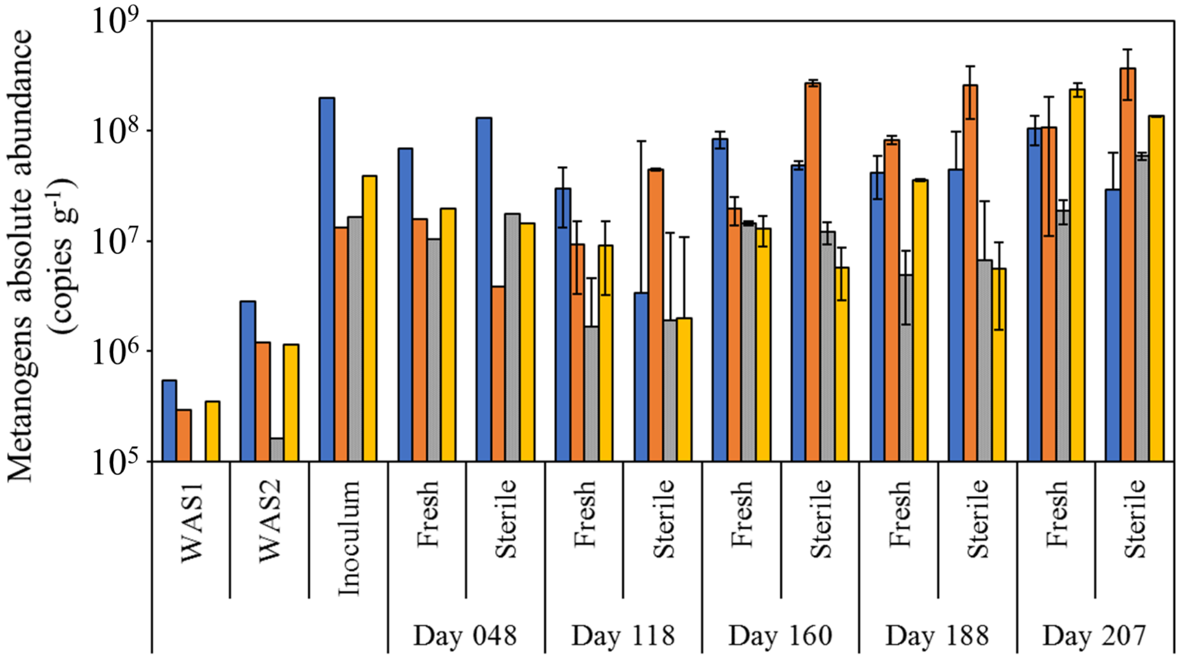

**Figure S8** Absolute abundance (copies $g^{-1}$) of the *Methanosaetaceae* (blue, ■), *Methanosarcinaceae* (orange, ■), *Methanobacteriales* (grey, ■) and *Methanomicrobiales* (yellow, ■) in the methanogenic community of the two batches waste activated sludge (WAS1 & 2), the inoculum and in the digesters fed fresh and sterile waste activated sludge on day 48, 118, 160, 188 and 207. Average values of the biological replicates (n=3) are presented, and the error bars represent standard deviations.
